# High level production and characterization of truncated human angiotensin converting enzyme 2 in *Nicotiana benthamiana* plant as a potential therapeutic target in COVID-19

**DOI:** 10.1101/2021.05.17.444533

**Authors:** Tarlan Mamedov, İrem Gürbüzaslan, Merve Ilgin, Damla Yuksel, Gunay Mammadova, Aykut Ozkul, Gulnara Hasanova

## Abstract

The COVID-19 pandemic, which is caused by SARS-CoV-2 has rapidly spread to more than 222 countries and has put global public health at high risk. The world urgently needs safe, a cost-effective SARS-CoV-2 coronavirus vaccine, therapeutic and antiviral drugs to combat the COVID-19. Angiotensin-converting enzyme 2 (ACE2), as a key receptor for SARS-CoV-2 infections, has been proposed as a potential therapeutic target in COVID-19 patients. In this study, we report high level production (about ∼0.75 g /kg leaf biomass) of glycosylated and non-glycosylated forms of recombinant human truncated ACE2 in *Nicotiana benthamiana* plant. The plant produced recombinant human truncated ACE2s successfully bind to the SARC-CoV-2 spike protein, but deglycosylated ACE2 binds more strongly than the glycosylated counterpart. Importantly, both deglycosylated and glycosylated forms of AEC2 stable at elevated temperatures for prolonged periods and demonstrated strong anti-SARS-CoV-2 activity *in vitro*. The IC_50_ values of glycosylated and deglycosylated AEC2 were 0.4 and 24 μg/ml, respectively, for the pre-entry infection, when incubated with 100TCID_50_ of SARS-CoV-2. Thus, plant produced truncated ACE2s are promising cost-effective and safe candidate as a potential therapeutic targets in the treatment of COVID-19 patients.

## Introduction

SARS-CoV-2 is a novel and highly pathogenic coronavirus, which has caused an outbreak in Wuhan city, China in 2019, and then soon spread nationwide and spilled over to other countries and the world, which resulted hundreds of thousands of deaths all over the world. Head of the United Nations has described this humanity’s worst crisis since World War II. Although several COVID-19 vaccines are currently available and a number of candidate vaccines are under development, only limited data are available on the effectiveness and safety of these vaccines. The world urgently needs effective and safe SARS-CoV-2 coronavirus vaccines, antiviral and therapeutic drugs to combat the pandemic that kills thousands of people every day. The development of therapeutic drugs can be a useful and alternative approach to suppress the virus entering and spread of the virus. Since angiotensin converting enzyme 2 (ACE2), the SARS-CoV-2 (COVID-19) receptor, is a critical molecule in the entry process of the virus into host tissue cells, it could be a potential therapeutic agent. ACE2 is a zinc containing metalloenzyme, present in most organs, attached to the cell membranes of cells in the lungs, heart, kidney, arteries and intestines^1^. ACE2 enzyme has multiple function and its primary function is to cleave the angiotensin I hormone into the vasoconstricting angiotensin II.ACE2 is a transmembrane protein, and serves as receptors for some coronaviruses such as SARS-CoV, SARS-CoV-2 and HCoV-NL63 ^2-7^. Similar to SARS-CoV, SARS-CoV-2 has been shown to bind to its functional receptor ACE2 via receptor binding domain (RBD) of SARS-CoV-2 spike protein as an initial step for entry into the cell^8,9^. It has been demonstrated that the binding affinity between ACE2 and RBD of SARS-CoV-2 is much stronger than that of SARS-CoV^9^, which can be logically explained the increased infectivity of SARS-CoV-2 versus SARS-CoV. It has been demonstrated that, ACE2 serves not only the entry receptor for SARS-CoV or SARS-CoV-2 but also can protects from lung injury^4,5,10-12^.Therefore, ACE2 has been proposed as potential therapeutic target to be used for SARS-CoV-2 infection^3,14^. Soluble ACE2 has also been described as a therapeutic candidate, which could neutralize the infection by acting as a decoy^15^.

Recombinant human ACE2 is also proposed as a novel treatment to improve pulmonary blood flow and oxygen saturation in piglets^16^. Based on reported pathological findings^17-19^, it has been shown that SARS-CoV-2 is associated with lung failure and acute respiratory distress syndrome. Pulmonary arterial hypertension (PH) is a devastating lung disease, which is characterized by high blood pressure in the pulmonary circulation^20^. From this point of view, the introduction of soluble recombinant human ACE2 into the human body has been proposed for the treatment of Acute respiratory distress syndrome (ARDS) and pulmonary arterial hypertension^21^. It should be noted that the Phase I (NCT00886353) and Phase II (NCT 01597635) clinical trials for recombinant human ACE2 have been successfully completed. The administration of soluble recombinant ECE2 has demonstrated safety and efficacy for the treatment of ARDS and without clinically significant changes in healthy people, as well as in patients with ARDS^22-23^. During the period of taking the drug, there were no serious adverse events no antibodies to recombinant human ACE2 detected^22^. Recently, it was shown that recombinant human ACE2 was significantly inhibited SARS-CoV-2 infection of Vero E6 cells^24^ and can neutralize SARS-CoV-2 infectivity in human kidney organoids^25^, Human Capillary and Kidney Organoids^24^. It was previously proposed that SARS-CoV may deregulate a lung protective pathway^4,10^. Thus, recombinant AEC2 might not only reduce lung injury, but also could block early the entry of SARS-CoV-2 infections in target cells. Therefore, the development of a cost-effective, safe, and functionally active recombinant ACE2 could be very important in the treatment of patients with COVID-19. Recent studies have shown plant expression systems as promising expression platforms for the rapid, safe and cost-effective production of various recombinant proteins. Plant expression systems have a number of advantages compared to other expression systems currently used, and have the ability to accumulate hundreds of milligrams of target protein per kilogram of biomass in less than a week. This system has been successfully used for rapid and cost-effective production of a variety of recombinant proteins such as vaccine candidates^26-30^ and complex mammalian proteins, enzymes such as furin and Factor IX^30^. Notable, using transplastomic technology human ACE2 was produced in plant chloroplasts^31^. It was demonstrated that the delivery of human ACE2 (fused with CTB) by oral gavage in mice resulted in increased circulating and retinal levels of ACE2 and reduced eye inflammation ^31^. Expression of ACE2 fused with the Fc region of human IgG1 in *Nicotiana benthamiana* plant using transient expression platform has been recently reported^32^. The expression level of ACE2 fusion protein was 100 mg/kg plant leaf. It was reported that this plant-produced fusion protein exhibited potent anti-SARS-CoV-2 activity *in vitro*.

ACE2 enzyme were also produced in various mammalian cells such as HEK293, CHO, insect cells etc. (https://www.mybiosource.com/search/Angiotensin-converting+enzyme+2). However, mammalian expression systems are extremely expensive and difficult to scale up. In addition, there is a risk of contamination of mammalian pathogens in recombinant proteins produced using the mammalian expression systems. Here we describe the engineering, expression and production at high level of truncated form of ACE2 in *N. benthamiana* plant using transient expression system, for the first time. The expression levels of both glycosylated and *in vivo* Endo H deglycosylated forms are ∼0.75 g/ kg leaf biomass. The purification yields of recombinant plant produced ACE2 protein glycosylated and deglycosylated forms calculated as∼0.4 or 0.5 g/kg leaf biomass, respectively. Expression of ACE2 proteins and purification procedures can be optimized to increase their purification yield. Our results demonstrated that the plant-produced truncated ACE2 proteins bind successfully to RBD of the SARC-CoV-2 spike protein and inhibit SARS-CoV-2 infection *in vitro*. In addition, our results demonstrate that both glycosylated and deglycosylated AEC2 proteins are stable at an elevated temperature for prolonged periods.

## Results

### Engineering and production, purification of recombinant ACE2s in *N. benthamiana* plants

We engineered and produced a truncated version of human ACE2 in *N. benthamiana* plant as described in Materials and Methods. To understand the role of glycosylation, we produced both glycosylated and non-glycosylated variants of ACE2 protein in *N. benthamiana* plant. Figure 1 demonstrates the confirmation of the production of glycosylated and non-glycosylated variants of ACE2 in *N. benthamiana* by western blot analysis. *N. benthamiana* leaf samples were harvested at different post infiltration days (dpi) and expression levels of glycosylated and non-glycosylated variants of ACE2 reached the maximum level at 6 dpi. For purification, a vacuum infiltration was used for large-scale production of glycosylated and non-glycosylated variants of ACE2. Glycosylated and deglycosylated variants of ACE2 were purified using HisPur™ Ni-NTA resin. The purification yields of recombinant plant produced glycosylated or deglycosylated forms were ∼0.4 or 0.5 g/kg leaf, respectively. The purity of glycosylated and deglycosylated variants of ACE2 enzyme was higher than 90% or 95%, for glycosylated or deglycosylated, respectively, as estimated based on SDS-PAGE and western blot analysis (Figure 2). Molecular masses were 80 or 90 kDa for deglycosylated or glycosylated AEC2, respectively (Figure 2).

**Figure 1.**
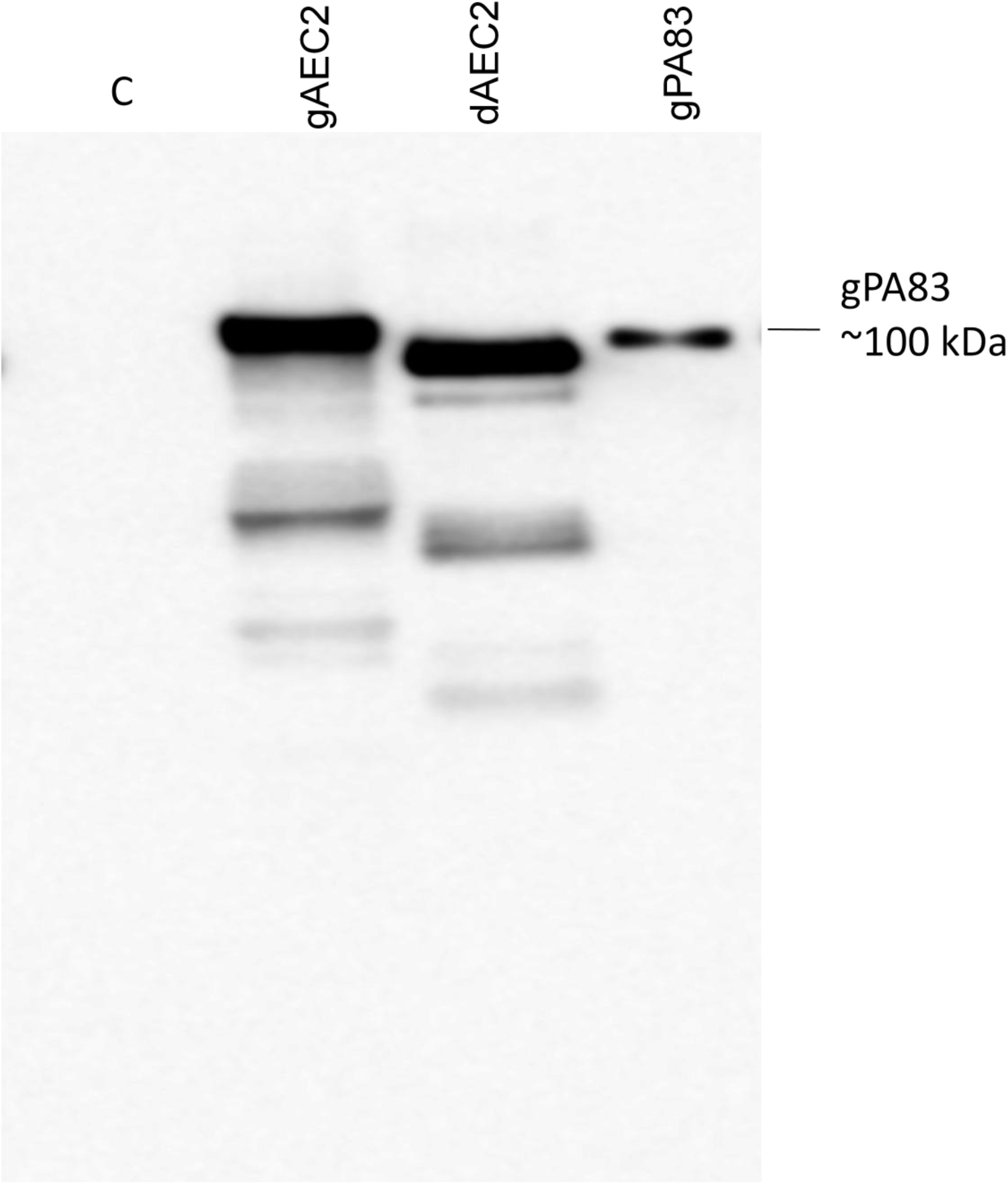
Western blot analysis of human ACE2s, produced *in N. benthamiana* plants. dACE2: ACE2 co-expressed with bacterial Endo H, produced in *N. benthamiana;* gACE2: Western blot analysis of human ACE2, produced in *N. benthamiana* plants; C-crude extract from non-infiltrated *N. benthamiana*; gPA83: PA83 of Bacillus anthracis,∼100 kDa, *N. benthamiana* leaf samples were harvested at 6 dpi.

**Figure 2.**
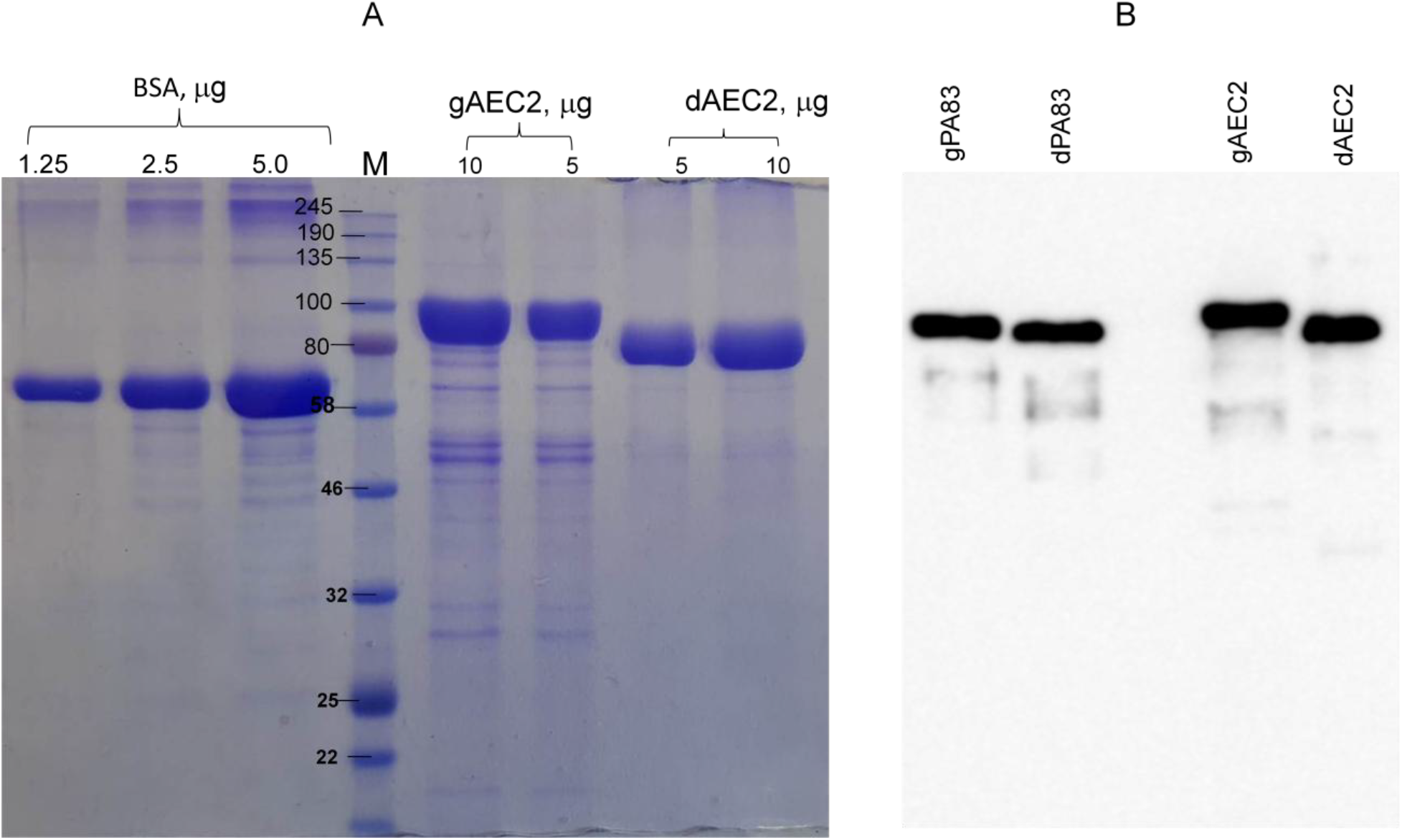
SDS-PAGE (A) and Western blot (B) analysis of plant produced, Ni-NTA resin purified glycosylated or deglycoslated ACE2 proteins. Glycosylated and deglycosylated plant produced ACE2 proteins were purified from *N. benthamiana* plant using HisPur™ Ni-NTA resin. gACE2: 5 or 10 μg purified glycosylated ACE2 protein was loaded into well; dACE2: 5 or 10 μg purified deglycosylated ACE2 proteins were loaded into well. BSA standards: 1.0, 2.5 and 5.0 μg BSA protein was loaded as a standard protein. B: membrane was probed with anti-His6 antibody. gPA83 (plant produced glycosylated Protective antigen of *Bacillus anthracis*, MM ∼100 kDa) and dPA83 (deglycosylated Protective antigen of Bacillus anthracis, MM ∼80 kDa) proteins were used as a standard. The image was taken using a highly sensitive GeneGnome XRQ Chemiluminescence imaging system.

### Binding affinity of plant produced recombinant ACE2 protein with spike protein

The binding activity of plant produced recombinant ACE2 protein was confirmed by measuring the binding activity of ACE2 with spike protein of SARC-CoV-2 as described in Materials and Methods. The results presented at Figure3 demonstrate that plant produced, glycosylated and non-glycosylated ACE2s successfully bind to commercial or plant produced spike proteins of SARC-CoV-2. Deglycosylated ACE2 variant binds to the deglycosylated plant-produced S-protein more strongly than the glycosylated counterparts.

**Figure 3.**
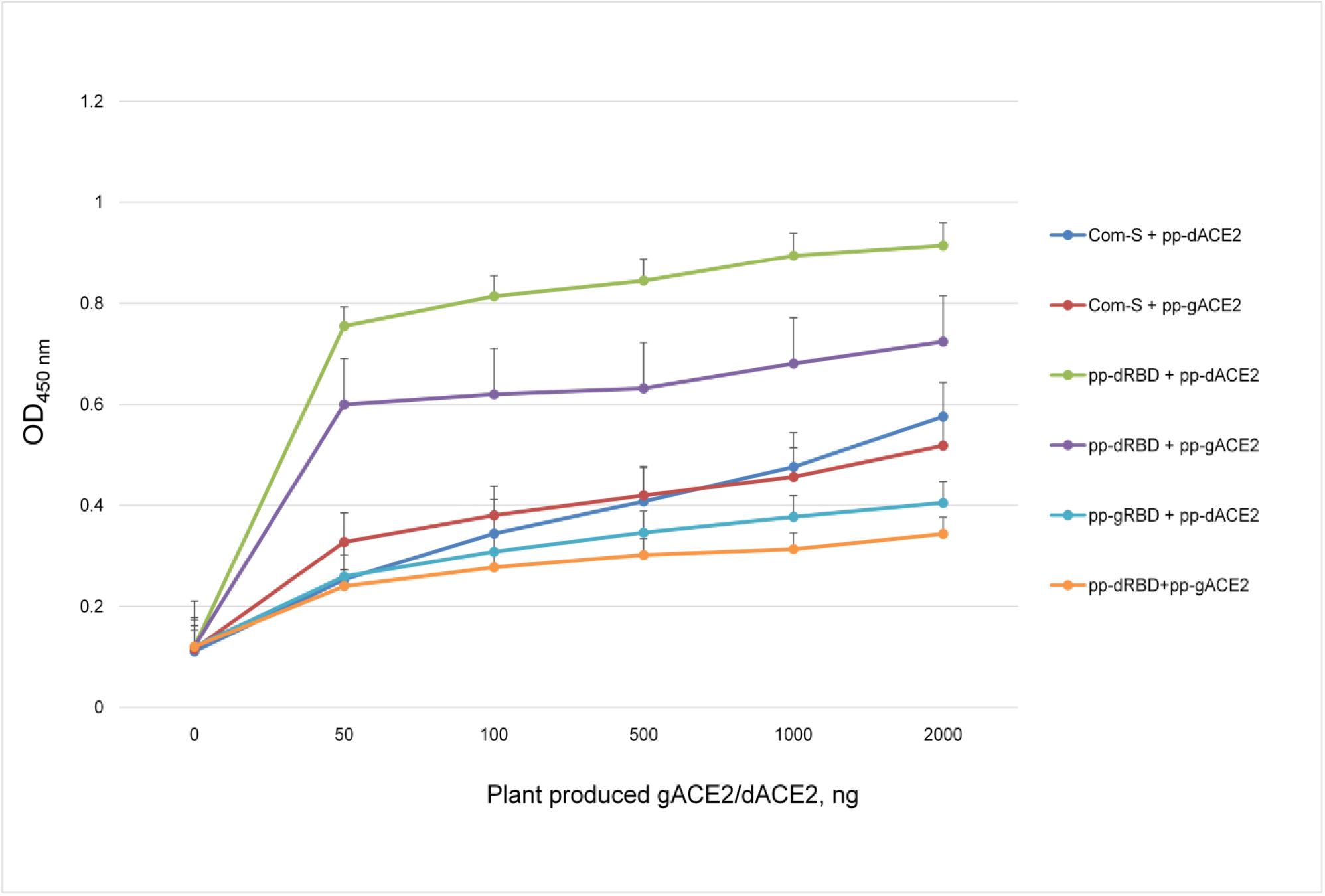
Binding activity of plant produced, glycosylated or deglycosylated variants of ACE2 with commercial or plant produced, glycosylated or deglycosylated forms of spike proteins (Flag tagged). Commercial or plant-produced spike protein was coated with an ELISA plate at a concentration of 200 ng /well. Different concentration of plant produced ACE2 (his tagged) was added. Purified anti-His Tag antibody (Cat. No. 652502, BioLegend) was used as a primary and mouse IgG used as secondary antibodies. SP (commercial): commercial Spike protein, active Recombinant 2019-nCoV Spike Protein, RBD, His Tag, produced in Baculovirus-Insect Cells, MBS2563882, Cat: MBS2563882); pp-gSP: plant produced glycosylated Spike protein; pp-dSP: plant produced deglycosylated Spike protein; pp-gACE2: plant produced glycosylated ACE2; pp-dACE2: plant produced Endo H in vivo glycosylated ACE2.

### Stability assessment of plant produced ACEs

The stability of plant produced glycosylated and *in vivo* deglycosylated forms of AEC2 were examined after incubation at 37°C for a prolonged time period: 24, 48, 96 and 144 hours (Figure 4). Analysis by SDS-PAGE showed that plant produced glycosylated ACE2 had almost no degradation at 37°C for 144 h and degradation of *in vivo* Endo H glycosylated ACE2 at the same condition was less than 5 %. These results demonstrate that plant produced, both glycosylated and deglycosylated AEC2s are stable at an elevated temperature for prolonged periods.

**Figure 4.**
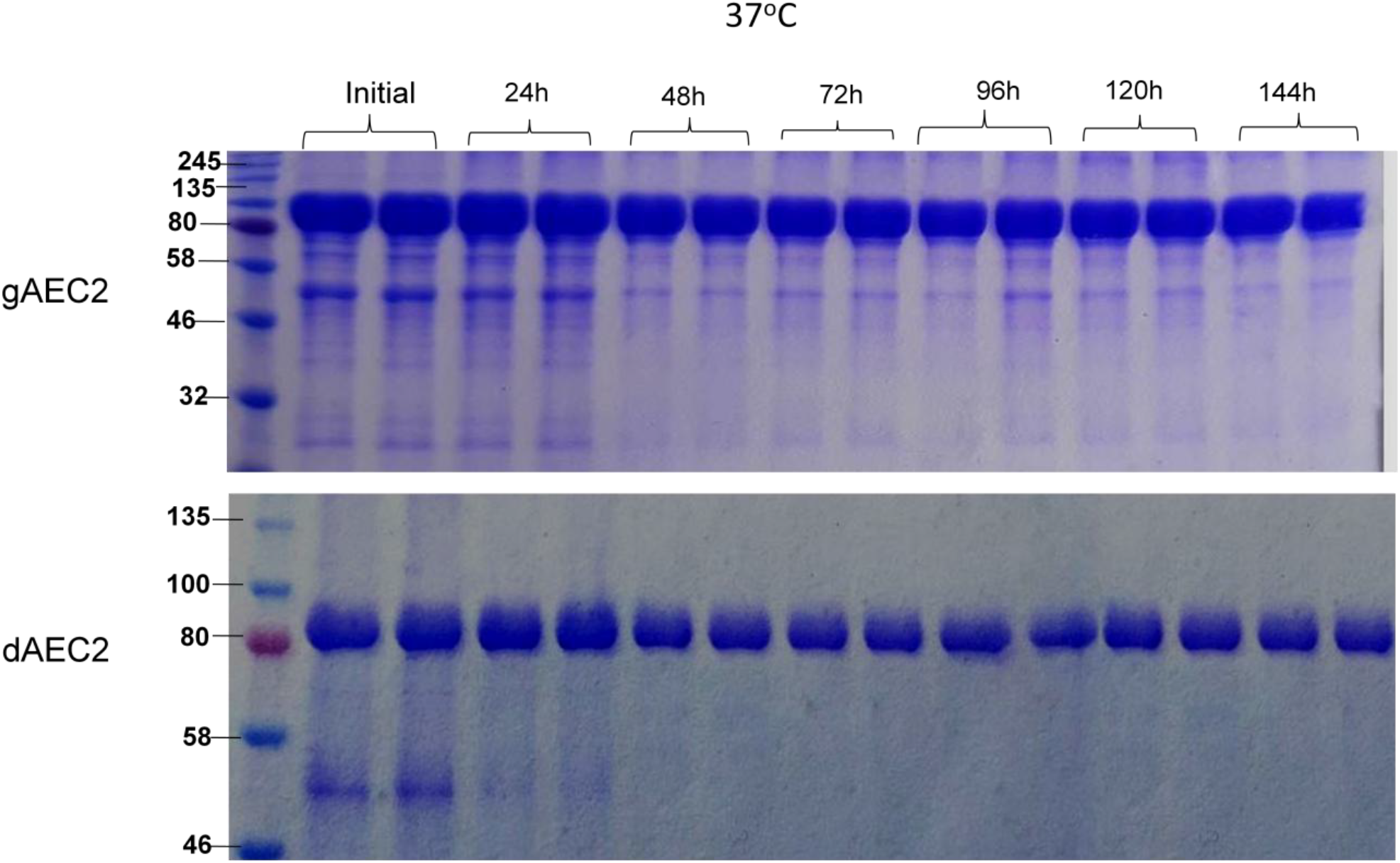
Stability assessment of plant produced glycosylated (A) and deglycosylated (B) ACE2 proteins. Plant produced, Ni-NTA resin column purified gACE2 or dACE variants were incubated at 37 °C for 24, 48, 72, 96 and 144 hours, and analyzed in SDS-PAGE. Lanes were loaded with ∼5.0 μg gACE2 (A) or d ACE2 (B). M: color prestained protein standard.

### Anti -SARS-CoV2 activity of plant produced ACE2s

Anti -SARS-CoV2 activity of plant produced glycosylated and deglycosylated forms were evaluated as described in Materials and Methods. Figure 5 demonstrate apparent activities of plant produced recombinant truncated glycosylated and deglycosylated ACE2 variants to RBDs plotted against IC50 of authentic SARS-CoV-2 neutralizationat the pre-infection phase. The half maximal inhibitory concentration (IC_50_) values for glycosylated and deglycosylated AEC2 were 0.4 and 24 μg/ml, respectively, when they mixed with 100TCID_50_ of SARS-CoV-2 (Figure 5).

**Figure 5.**
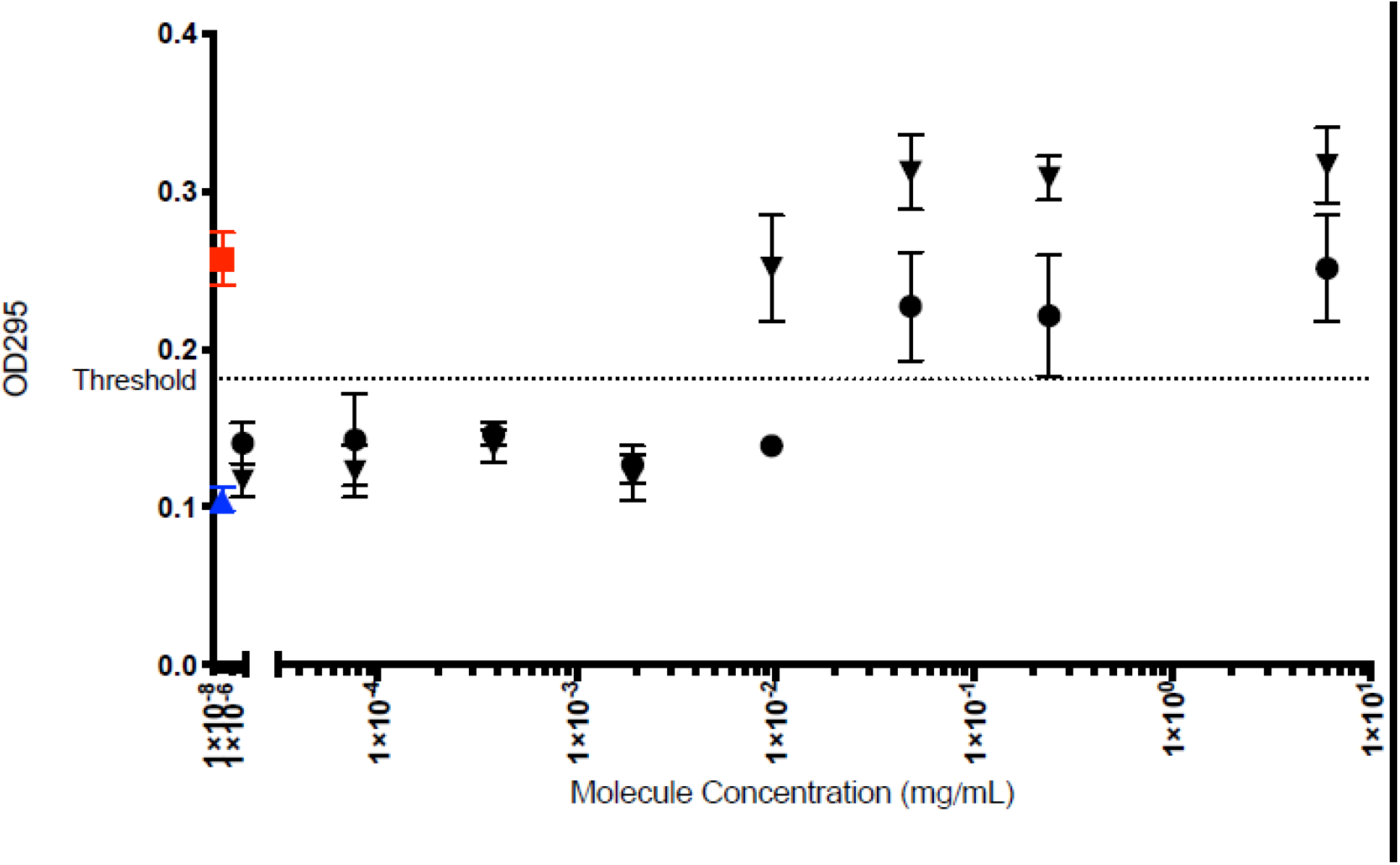
Apparent activities of two distinct ACE2 derivatives produced in plants to RBDs plotted against IC50 of authentic SARS-CoV-2 neutralization.

## Discussion

The new coronavirus COVID-19 disease is currently responsible for the pandemic, created huge global human health crisis with significant negative impacts on health and economy worldwide. Given the high rates of morbidity and mortality associated with COVID-19, there is an urgent demand for the development of effective, safe and affordable therapeutics, vaccines and inhibitors to control the epidemic. A number of studies performed with SARS-CoV and SARC-CoV-2 have been demonstrated the ACE2 enzyme as a potential therapeutic target and alternative treatment option in COVID-19 patients. Recombinant human ACE2, developed by APEIRON Biologics (https://www.apeiron-biologics.com/tag/ace2/), is currently in pilot clinical trials in China. Thus, the high-level production of safe and affordable recombinant human ACE2 with high anti SARS-CoV-2 activity would be a challenge task. Therefore, the aim of this study is to produce an affordable, safe and functional active recombinant human ACE2 using the plant transient expression system for use in the treatment of COVID-19. Plant expression systems is promising expression platform for cost-effective, fast and safe production of a variety of recombinant proteins, vaccines, antibodies, therapeutic proteins and enzymes. Plant expression systems have several advantages over other expression systems currently in use, including rapid and high production with the ability to accumulate grams of target protein per kilogram of biomass in less than a week. This system have been successfully used for production functional active complex proteins such as full length Pfs48/45 of *Plasmodium falciparum*^30^, human Furin, Factor IX^30^, glycohormone erythropoietin^33^, latent transforming growth factor-beta (TGF-beta)^34^, and also PA83 of *Bacillus anthracis*^27, 28^, HIV gp140 and other viral glycoproteins^35,36^. The goal this study is to develop safe, cost effective plant produced ACE2 enzyme with anti-SARS-CoV-2 activities for use in the treatment of COVID-19.

ACE2 is zinc-metalloproteinase type 1 transmembrane protein with molecular mass of 120 kDa and composed of 805 amino acids. 3D structure analysis of ACE2 protein was revealed that enzyme molecule contains a signal peptide sequence (1-17 aa) and extracellular sequence (18-740) which contains the active carboxy peptidase domain, a transmembrane domain (aa 741– 761) and a cytoplasmic domain (aa 762–805). The ACE2 enzyme has been intensively studied since 2002 as it was identified as a cellular receptor for the SARS-CoV, HCoV-NL63 and SARS-CoV-2 coronaviruses.

In this study, we engineered and expressed a truncated form of human ACE2 gene in *N. benthamiana* plant, for the first time. Spike-glycoprotein of SARC-CoV-2 has 22 potential N-glycosylation sites. Human ACE2 is glycoprotein, which has 7 potential N-glycosylated sites. The virus (SARS-Cov-2) and receptor (ACE2) binding affinity on the surface of human cells could be critical step in viral entry into the susceptible cells. To understand the role of N-glycosylation, we produced both glycosylated and non-glycosylated variants of ACE2 protein in *N. benthamiana* plant. Deglycosylated ACE2variant was produced using the *in vivo* deglycosylation strategy, by co-expression of human ACE2 with bacterial Endo H, which we recently developed^28^. The expression level of glycosylated and deglycosylated variants was about ∼0.75g/kg plant biomass. Notable, the production of soluble human ACE2 as a fusion protein with immunoglobulin Fc domain has been recently reported ^32^.The expression level of ACE2-Fc fusion reported was 100 mg/kg plant leaf. The expression levels of the truncated ACEs developed in this study are 7.5 times higher than the ACE2-Fc fusion. Both glycosylated and deglycosylated truncated ACE2 variants were purified using Ni-column affinity chromatography. The purification yield of glycosylated and deglycosylated ACE2 variants were ∼0.4 and ∼0.5 g/kg plant leaf. The purities of glycosylated or deglycosylated AEC2s were 90% or 95 %, respectively.

Our results showed that plant produced glycosylated and non-glycosylated ACE2 variants successfully bind to RBD of SARC-CoV-2 spike protein. However, the deglycosylated ACE2 variant binds to the deglycosylated plant-produced S-protein more strongly than the glycosylated counterparts. Moreover, the plant produced glycosylated and non-glycosylated ACE2 variants exhibited potent anti-SARS-CoV-2 activity *in vitro*. However, the IC_50_ values of glycosylated ACE2 (0.4 μg/ml) was 60 -fold lower compared to deglycosylated form of ACE2 (24 μg/ml), for the pre-entry infection, when culture medium was mixed 100TCID_50_ of SARS-CoV-2. Notable, IC_50_ values of ACE2-Fc fusion protein at the pre-entry stage was reported as 94.66 μg/ml at 25TCID_50_ ^32^.Thus, the inhibition efficacy of plant produced glycosylated, truncated form ∼947 fold higher than ACE2-Fc.

Generally, it has been supposed that soluble form of ACE2 in excessive forms, may negatively affect the virus entering and spreading^37^. Based on above descriptions plant produced recombinant ACE2 could be as a potential therapeutic target in COVID-19 patients to slow down the virus entering, spread of the virus and protect the lung from injury. Like other respiratory diseases, COVID-19 can cause permanent damage the lungs, heart, and other organs. A possible explanation is the blocking of the binding domain of the ACE2 receptor (RBD) by SARC-CoV-2. Therefore, recombinant ACE2 could be a promising bio-mimicking molecule facilitating to attenuate and/or prevent COVİD-19 related cellular injury. Thus, development of safe, functional active and a cost-effective production of recombinant ACE2 could be very useful for the treatment of COVID-19 patients. Collectively, all above findings demonstrate that plant produced glycosylated and deglycosylated forms of truncated ACE2 could be a potential therapeutic agent against SARS-CoV-2 to slow down the virus entering, spread of the virus and protect the lung from injury.

### Conclusion

A number of studies have been shown recombinantACE2 as a potential therapeutic target in COVID-19 patients. At this point, the development and production of recombinant ACE2 protein at high levels with high anti SARS-CoV-2 activity could be a challenging task. In this study, we show that recombinant ACE2 that exhibit a potent anti-SARS-CoV-2 activity with the IC_50_ values of 0.4 μg/ml, can be produced rapidly, at high level (∼ 0.75 g/kg plant leaf) in *N. benthamiana* plant using plant transient expression system. These findings demonstrate plant produced ACEs as a cost effective, safe and promising therapeutic target for the treatment of COVID-19 patients.

## Materials and Methods

### Cloning, expression, and screening of recombinant ACE2 in *N. benthamiana* plants

The sequences of ACE2 was optimized for expression in *N. benthamiana* plants and synthesized de-novo. The constructed ACE2 gene was inserted into the pEAQ binary expression vector ^38^ to obtain pEAQ-ACE2. pEAQ-ACE2 plasmid was introduced into the Agrobacterium tumefaciens strain AGL1. AGL1 carrying the pEAQ-ACE2 plasmid was then infiltrated into 6-7-week old *N. benthamiana* plants. To produce deglycosylated ACE2, ACE2 gene was *in vivo* co-expressed with Endo H^28^. To confirm the expression of His6 tagged ACE2 protein variants, a leaf tissue was harvested at 6 dpi (day post infiltration) and homogenized in three volumes of extraction buffer (20 mM sodium phosphate, 150 mM sodium chloride, pH 7.4) as described previously^29^.

### Purification of recombinant ACE2 from *N. benthamiana* plants

To produce the AEC-2 protein (both glycosylated and deglycosylated variants) in *N. benthamiana*, plants were infiltrated with ACE2 (glycosylated) or ACE2 + Endo H (deglycosylated) genes and harvested at 6 dpi. For purification, twenty grams of frozen plant leaves from each variant infiltrated with AEC-2 gene were ground in extraction buffer with 3 times volume of plant weight and the extract was centrifugated for 20 minutes at 4°C at 13,000 g. The supernatant was loaded onto a disposable polypropylene column (Pierce) with 1 ml HisPur™ Ni-NTA resin equilibrated with 10 column volume binding buffer, by gravity-flow chromatography. The column was washed with 10-15 column volumes (CV) of wash buffer until reaching to the baseline. Proteins were eluted with 10 CV of elution buffer. Elution fractions were collected as 0.5 ml/eppendorf and protein concentrations in the eluted fractions were measured by BioDrop. According to the concentration, the combined fractions were concentrated, and buffer exchanged against PBS with a 10K MWCO Millipore concentrator (Cat No: UFC801096, Merck) to a final volume of 1.2 ml. The concentrated protein was stored at – 80°C until use.

### Study of binding activity of plant produced recombinant ACE2 protein with commercial or plant produced spike protein

Binding activity of plant produced recombinant ACE2 protein with commercial or plant produced spike proteins was performed by ELISA. Briefly a 96-well plate (Greiner Bio-One GmbH, Germany) was coated with 100 ng of plant produced^39^ or commercial available Spike protein of SARS-CoV-2 (RBD, His Tag, Arg319-Phe541, MM∼ 25 kDa, MBS2563882,MyBioSource, USA) in 100 mM carbonate buffer for overnight. Next day wells were blocked with blocking buffer for 2 h at room temperature. After blocking various concentrations of plant produced glycosylated and deglycosylated ACE2 proteins (100 −2000 ng) were added into wells and incubated for 2 h at 37 °C. After 2 h, anti-His tag mouse mAb was added into each wells. The plate was washed three times with blocking solution (200 μl/well). After washing, wells were incubated with anti-mouse HRP -IgG antibody (BioLegend). The plate was washed three times with washing solution (200 μl/well for 5 minute). 200 μl of substrate solution (Sigma) was added to each well. Afterwards plate was incubated in the dark, for 30 minutes at RT. After the incubation period, the plate was read at 450 nm on a multi well plate reader.

### Stability assessments of different variants of ACE2

Stability assessments of different variants of ACE2 were performed using similar procedure as described previously^28,30^. Plant produced glycosylated and deglycosylated variants of ACE2 were diluted to 1,0 mg/mL with PBS, and were aliquoted into low-binding tubes. Proteins were then incubated at 37 °C for 48, and 96 hours. After incubation, samples were analyzed by SDS-PAGE. For SDS-PAGE analysis, the samples were mixed with SDS loading dye (5X) and stored at −20 °C until use. All samples were then run on SDS-PAGE. The degradation of ACE2 variants were quantified using highly sensitive Gene Tools software (Syngene Bioimaging, UK) and ImageJsoftware (https://imagej.nih.gov/ij).

### SDS-PAGE and western blot analysis

SDS-PAGE and western blot analysis of plant produced glycosylated and non-glycosylated variants of ACE2 were performed as described previously^30^. The His tagged ACE2 variants were detected using anti-His mAb (Cat. No. 652505, BioLegend).

### Anti -SARS-CoV2 activity of plant produced ACE2s

Anti SARS-CoV2 activity of plant produced ACE2s was monitored in vitro. The neutralizing ability of glycosylated and deglycosylated ACE2 variants was analyzed at different incubation periods. dACE2 and gACE2 (3.055 and 2.542 mg/mL, respectively) was 5-fold diluted in high glucose DMEM in a U-bottomed plate. After combining with equal volume (100 µL) 100TCID50 of SARC-CoV-2 (10^4,25^/0.1 mL), the mixtures were incubated at room temperature for 30 minutes. A total of 150 µL incubated mixture was then inoculated on Vero E6 Cells grown in a 96-well flat-bottomed tissue culture plate (Greiner, Germany). The highest concentrations of deglycosylated ACE 2 and glycosylated ACE2 without the virus was involved as a toxicity control, and serum-free High Glucose DMEM was added to each plate as a cell control. A total of 75 µL of 100TCID_50_ SARS-CoV2 Ank1 virus was also used as virus control. All tests were performed in a quadruplicate. The plates were incubated at 37°C in a humidified incubator with a 5% CO_2_ atmosphere until virus control wells had adequate cytopathic effect (CPE) readings. The test was evaluated when the virus control wells showed 100% cytopathogenic effect (CPE) at daily microscopy. To do precise calculations based on OD values, cells were fixed with 10% formaldehyde for 30 minutes and subsequently stained with crystal violet (CV - 0.075% in ethanol) for 20 minutes. The dye washed away by repeated washing and retained CV was released by adding 100 µL ethanol (70%). Ten minutes after, the plate was read on ELISA reader using 295 nm filter (Multiskan Plus, MKII, Finland). The 50% effective dose (EC_50_) was calculated as described elsewhere^40^.

## Acknowledgments

The authors are grateful to Dr. George P. Lomonossoff (John Innes Centre, Biological Chemistry Department) and Plant Bioscience Limited for kindly providing pEAQ binary expression vector. This study was funded by Akdeniz University.

## Author Contributions

T.M. Conceived the study; T.M., A.O designed the experiments; I.G., M.I., D.Y. performed the experiments. T.M., G.M., G.H., A.O. analyzed the data. T.M., G.M., G.H. contributed to writing the manuscript.

All authors reviewed the manuscript.

